# Organotypic Timelapse recording with Transcriptomic Readout (OTTR) links cell behaviour to cell identity in human tissues

**DOI:** 10.1101/2025.07.28.667119

**Authors:** Elin Vinsland, Camiel Mannens, David Fernández-García, Jokubas Janusauskas, Ivana Kapustová, Lijuan Hu, Peter Lönnerberg, Xiaofei Li, Xiaoling He, Roger Barker, Oscar Persson, Erik Sundström, Sten Linnarsson

## Abstract

Linking dynamic cellular behaviour to molecular states in intact human tissue remains challenging because during live imaging only limited molecular information can be captured while high-dimensional molecular measurements are destructive. Here we describe Organotypic Timelapse recording with Transcriptomic Readout (OTTR), which integrates week-long live imaging of sparsely labelled organotypic slice cultures with highly multiplexed *in situ* spatial transcriptomics. We applied OTTR to primary human glioblastoma and fetal cortical tissues. Using sparse labelling, we tracked the migration, proliferation, and lineage of tens of thousands of individual cells per sample. Following live imaging, precision resectioning and alignment allowed us to perform spatial transcriptomics on the very same tissue, thereby preserving the link between dynamic cell behaviours and transcriptomic states. We used OTTR to quantify cell-type specific migration patterns, lineage trees and the behaviour of cells near vasculature. OTTR provides a powerful, broadly applicable method for investigating the complex interplay between cell behaviour and molecular state in human tissues.

## INTRODUCTION

The functions of biological tissues are the result of coordinated cell behaviour driven by changes in cell molecular states. To understand tissue function, therefore, requires simultaneous monitoring of both dynamic cell behaviour — e.g. migration, cell division and interaction— and the underlying changes in cellular states. For example, in the developing human cerebral cortex, progenitor radial glia divide asymmetrically and their offspring (future excitatory neurons) migrate along the radial processes to settle in the correct layer; simultaneously, inhibitory neuron progenitors migrate perpendicularly to the radial processes while differentiating into many specialised interneurons. This complex dynamic process is driven by carefully regulated changes in transcriptional states. In brain cancer, these same processes are co-opted but transformed, leading to aberrant proliferation, differentiation, and interaction with surrounding normal tissue^1,2^.

Our understanding of these processes has been greatly aided by new technologies. For instance, single-cell RNA sequencing (scRNA-seq) has been used to describe how glioblastoma tumour cells resemble neurodevelopmental cell types^3^, and spatial transcriptomic methods have revealed microenvironmental drivers of glioblastoma organisation^4,5^. Recent advances in lineage tracing with genetic barcodes has also allowed the mapping of lineage progression and clonal relationships in mouse brain development^6^, as well as cellular plasticity and clonal drug resistance in glioblastoma neurospheres^7^.

However, these technologies capture only a temporal snapshot of a tissue or lose spatial information upon cell dissociation for sequencing. Yet spatial context is crucial both during normal brain development and brain tumour growth. Currently, no method exists that captures both the dynamic behaviour of many cell types, and their molecular and spatial information in the same tissue. To address this gap, we developed OTTR, a method that combines long-term live imaging of organotypic slice cultures, with highly multiplexed mRNA *in situ* analysis (10X Genomics Xenium, hereafter ‘spatial transcriptomics’), to capture both dynamic cell-type behaviour and single-cell transcriptomic signatures in the same tissue (Fig. 1a).

**Figure 1:**
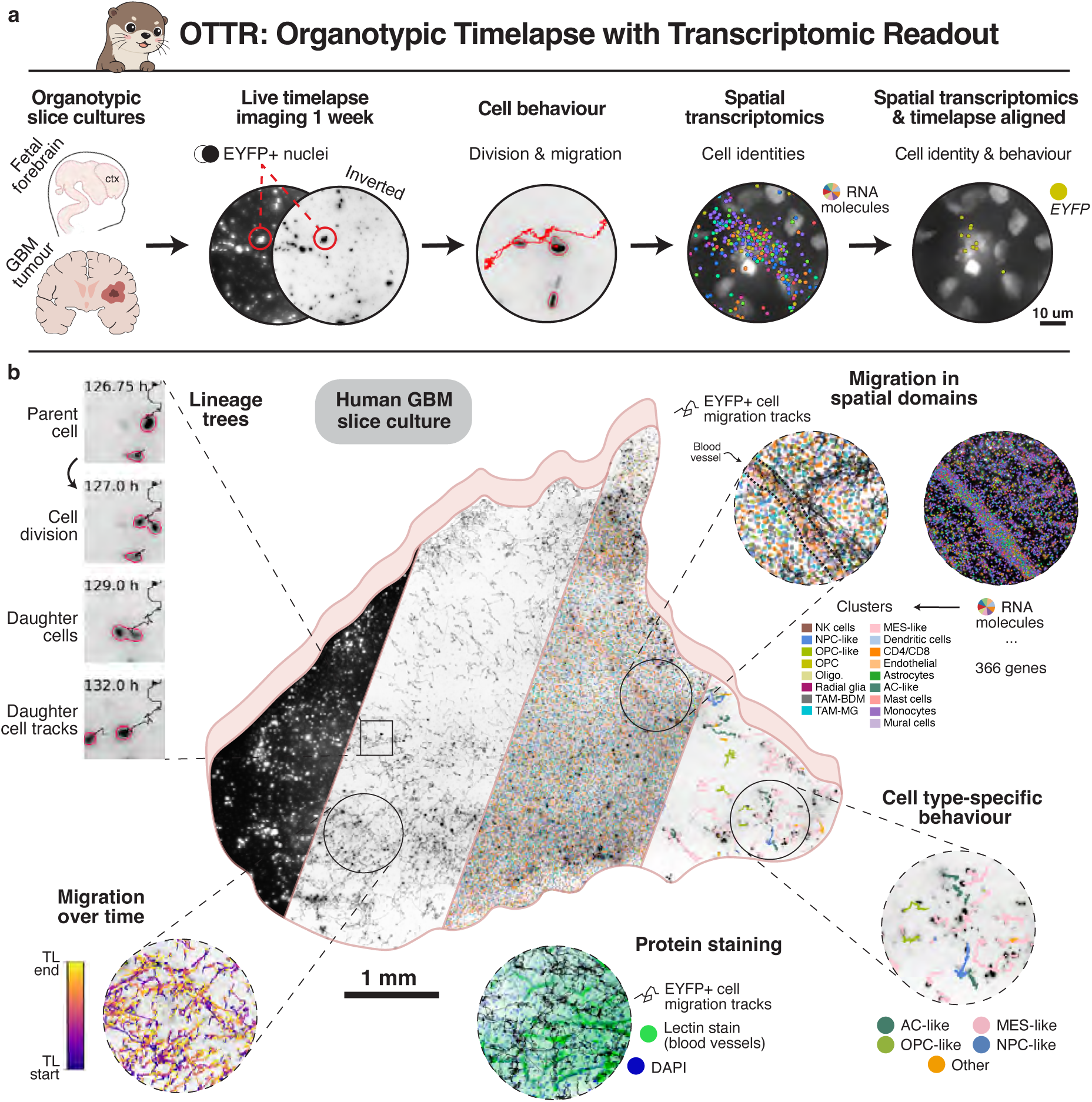
OTTR - Organotypic Timelapse with Transcriptomic Readout. **a)** Overview of experimental workflow and the data acquired. Live timelapse imaging was performed to track the behaviour of EYFP-fluorescent cell nuclei, followed by spatial transcriptomics and alignment of tracked cells with their molecular identity. ctx; cortex, GBM; glioblastoma. **b)** Left to right - the type of data collected with our method, using timelapse imaging data alone, spatial transcriptomic data alone, combined timelapse cell migration tracks with spatial transcriptomic clusters, and tracks of cells that were also aligned to the spatial transcriptomic data, linking cell behaviour to their molecular identity. TL; timelapse.

To facilitate cell tracking, we used sparse lentivirus labelling with a fluorescent reporter. Tens of thousands of cells per slice were labelled and continuously imaged on a fully automated microfluidic culturing and imaging system for up to a week. Afterwards, the slice was re-embedded in gelatin and re-sectioned on a cryostat. We used precise procedures to ensure that the cryosections would be aligned with the imaged part of the slice.

Finally, the cryosections were subjected to spatial transcriptomics and aligned to the last timelapse frame. The resulting dataset was analysed on three levels, by (1) quantifying proliferation, lineage trees and migration patterns using the timelapse data alone (Fig. 1b), (2) quantifying cell behaviour across spatial niches using the timelapse and spatial data aligned on the tissue level — e.g. cell migration around vasculature (Fig. 1c), and (3) performing cell type-specific analyses of individual tracked cells also aligned to the spatial data (Fig. 1d, Extended Data Fig. 1).

By combining long-term live imaging and highly multiplexed spatial transcriptomics, our method captures cell-type specific behaviours in primary human tissue. This allows us to study biology previously inaccessible in human tissues, where advances in *in vitro* and *ex vivo* human tissue models are essential to progress in the field^10^. In this paper, we provide comprehensive instructions for building the necessary instrumentation and performing complete end-to-end experiments, including detailed laboratory protocols and computational procedures. We also present examples of the kinds of analyses possible on human primary tissues, along with the complete raw data generated throughout.

## RESULTS

### Explant cultures and live imaging

First, we generated organotypic slice cultures, labelled cells using virally delivered ubiquitous Enhanced yellow fluorescent protein (EYFP)-expression, and imaged for one week on an automated microfluidic culturing and imaging system. Patient high-grade glioblastoma tumours (using WHO 2021 classification) were sectioned to 300 μm using a vibratome (Methods). Human fetal cortices up until PCW 10 were thin enough such that vibratome sectioning was unnecessary. Instead, we cut them into strips along the dorso-ventral axis and directly placed en face, ventricular side down, on PTFE culturing membranes (Fig. 2a, Extended data Fig. 2a-c).

**Figure 2:**
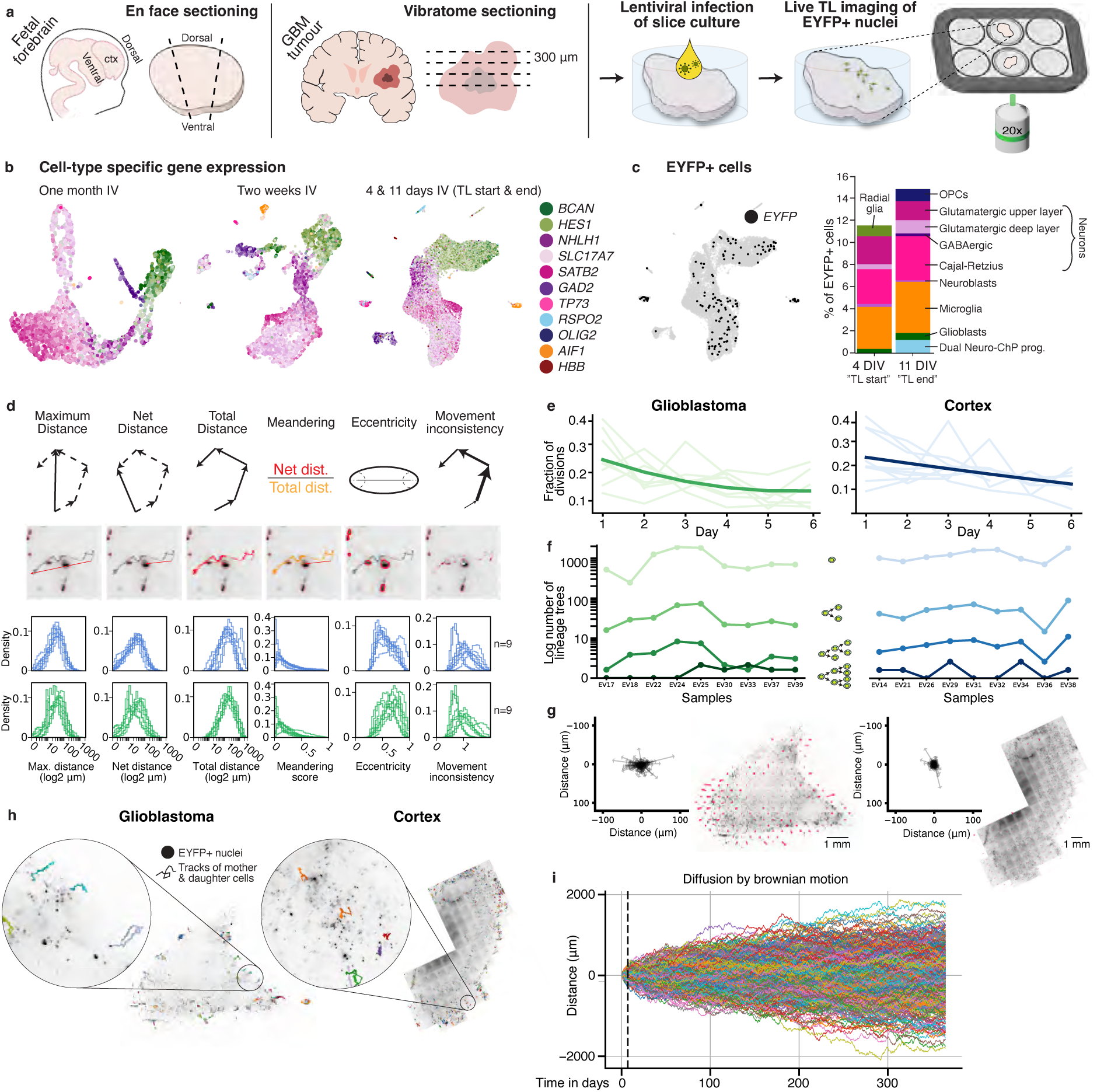
Cell tracking in cultured explants from fetal cortex and patient-derived glioblastoma tumours. **a)** Collection and processing of samples. Infection with a H2B-EYFP carrying lentivirus allows visual tracking of individual cells in *ex vivo* slice cultures. **b)** scRNA-seq UMAPs of cultured cortex explants at one month, two weeks and four and 11 days *in vitro*, with cells coloured by the expression of the gene they express the most highly among those in the key. IV, *in vitro*. **c)** scRNA-seq UMAP coloured by cells expressing at least one molecule of EYFP, and stacked bar charts showing the percentage of EYFP-positive cell types at four and 11 days *in vitro*. **d)** Distributions of movement characteristics per timelapse recording of cortex (blue) and glioblastoma (green) samples. From left to right the maximum distance from start position, net movement from start position, total distance travelled, fraction of net distance over the total distance travelled (meander), eccentricity (based on focal points of ellipse) of the nuclei and average deviation from average movement speed (movement inconsistency) are shown. **e)** Average fraction of cell divisions identified during each 24 hour interval of cell tracking. Individual samples are shown in the background. **f)** Quantification of the number of divisions occurring in each lineage tree for each sample. **g)** Mean movement of cells within 20-by-20 hexagonal grid for two representative samples from the cortex and glioblastoma, arrows are scaled to 4 times the actual movement. **h)** Lineage trees derived from tracking cell divisions of EYFP+ cells, where separate tracks of the same tree are shown in different shades of the same colour. EYFP+ nuclei are black on a background of the whole tissue. **i)** Predicted diffusion of tumour cells into the surrounding tissue over the course of a year. The median diffusion coefficient across all tracked glioblastoma samples was used. Dashed line indicates the end of the timelapse experiment.

To assess tissue health with our culturing method, we performed scRNA-seq analysis of cortical explants using adjacent slices from the same samples subjected to timelapse imaging. We cultured these control slices for four days *in vitro* (mimicking the start of a live imaging experiment after lentiviral infection and expression of EYFP), 11 days *in vitro* (corresponding to the end of one week of live imaging), and for two weeks and one month *in vitro*. At each of the four timepoints, we analysed cells that passed our quality control (Methods) (Extended data Fig. 2d). All four timepoints showed the expected cell types found in developing forebrain, such as cycling neural progenitors (*BCAN*+, *HES1*+), neuroblasts (*NHLH1*+), glutamatergic deep-layer (*SLC18A7*+), and upper-layer (*SATB2*+) neurons, Cajal-Retzius cells (*TP73*+), GABAergic neurons (*GAD2*+), oligodendrocyte progenitor cells (*OLIG2*+), and microglia (*AIF1*+). Two samples also included *RSPO2*+ neural progenitors, potentially due to anatomical differences from slicing the forebrain (Fig. 2b, Extended data Fig. 2e-g). While we only performed one scRNA-seq experiment at four and 11 days *in vitro*, here we observed fewer radial glia, fewer glioblasts, but more neuroblasts after one week of culturing (Extended data Fig. 2h-i).

Importantly, scRNA-seq analysis of explant cultures confirmed that *EYFP*+ cells were represented across all cell types (Fig. 2c), indicating that our lentivirus had transduced cells ubiquitously as intended. 0.5% and 0.85% of cells captured with scRNA-seq were *EYFP*+ at four and 11 days *in vitro*, respectively (Extended data Fig. 2j). Typically, we captured one to two *EYFP* mRNA molecules in *EYFP*+ cells (Extended data Fig. 2k).

During live imaging, we cultured tissue slices in a temperature and gas-controlled stagetop incubator (OKOlabs, Extended data Fig. 3). We used an automated microfluidic system to regularly replenish culture media, thereby avoiding sudden changes in nutrient availability and gas composition. Importantly, it also obviated any tissue handling during imaging, minimising the risk of tissue misalignment. Imaging was performed on a fully automated epifluorescence microscope (Nikon TiE2 Eclipse; Table S2). Following data processing, cells were tracked using a Bayesian tracking algorithm (btrack^11^). We report a total of 18 experiments (9 fetal cortex, 9 glioblastoma). On average, we tracked 29,550 EYFP+ cells per fetal cortex (295,495 total), and 11,274 per glioblastoma slice (225,470 total).

We calculated dynamic cell characteristics based on individual migration tracks and images^12^, including distance measures, nuclei eccentricity and movement inconsistency (Fig. 2d). These measures where highly consistent between imaging runs and tissue samples (Fig. 2d, bottom), but with some variation in movement inconsistency in both fetal cortex and glioblastoma as well as eccentricity in glioblastoma.

To reveal lineage relationships, we identified cell divisions (btrack; Methods; Video S1) and followed cells across divisions to create branched tracks. The rate of proliferation was higher in fetal cortex relative to glioblastoma, decreasing slowly over time (Fig. 2e). While most cells did not undergo mitosis during live imaging, nearly all sections contained at least several trees consisting of two or more cell divisions (Fig. 2e-f).

Cell tracking also allowed us to link cell division and lineage trees to cell migration, and to investigate local concerted migration (Fig. 2g-h, Video S1) across both glioblastoma and cortex samples. Interestingly, we found that cells in glioblastoma slices were more likely to exhibit locally concerted movement than in fetal cortex slices (Fig. 2g; Extended data Fig. 5). In part, this difference can be explained by the preparation of fetal cortex slices, which were positioned with the subventricular zone towards the bottom of the plate and most movement of neural progenitor cells and their progeny occurs radially^13^. In addition, we observed a greater tendency among glioblastoma cells to migrate away from the tissue centre compared to fetal cortex (Extended data Fig 4b).

Next, we asked to what extent undirected movement of glioblastoma cells could explain infiltration of tumour cells into the surrounding tissue. Assuming Brownian motion of tumour cells, we calculated the diffusion coefficient for all tumour samples and used the median value to model cellular migration (Fig. 2i; Extended data Fig. 4a). Based on their behaviour in tissue explants *ex vivo*, over the course of a year, most cells would not be expected to migrate more than 2 mm away from the tumour edge by undirected motion alone. Thus, directed migration, e.g. along white matter tracts or blood vessels, is required to explain the wide dissemination of glioblastoma tumour cells in patient brains.

### Resectioning of slice cultures and subsequent spatial transcriptomic analysis

Next, our aim was to determine the identities of the cells that were tracked during live imaging. For that, we established a protocol to resection the 300 μm cultured tissue slice into 12 μm sections for spatial transcriptomic analysis (full protocol in Extended data Fig. 6 and Methods). Briefly, the tissue slice was fixed and embedded in gelatin inside the culturing insert, excised, flipped vertically, and sandwiched between two flat slabs of gelatin on a cryostat chuck (Fig. 3a). The purpose of this procedure was to acquire cryosections starting from the bottom of the cultured tissue because live imaging was performed from this side with an inverted microscope, with around 60 μm depth of imaging into the tissue on the Z-plane. Furthermore, the angle of sectioning against the tissue en face needed to be as aligned and flat as possible, for later computational realignment between the spatial data from the 12 μm cryosections and the data from live imaging (more in Fig. 4): to fit a single cryosection entirely within the imaging volume across a centimetre slice required perfect flatness and less than 0.3 degrees of tilt.

**Figure 3:**
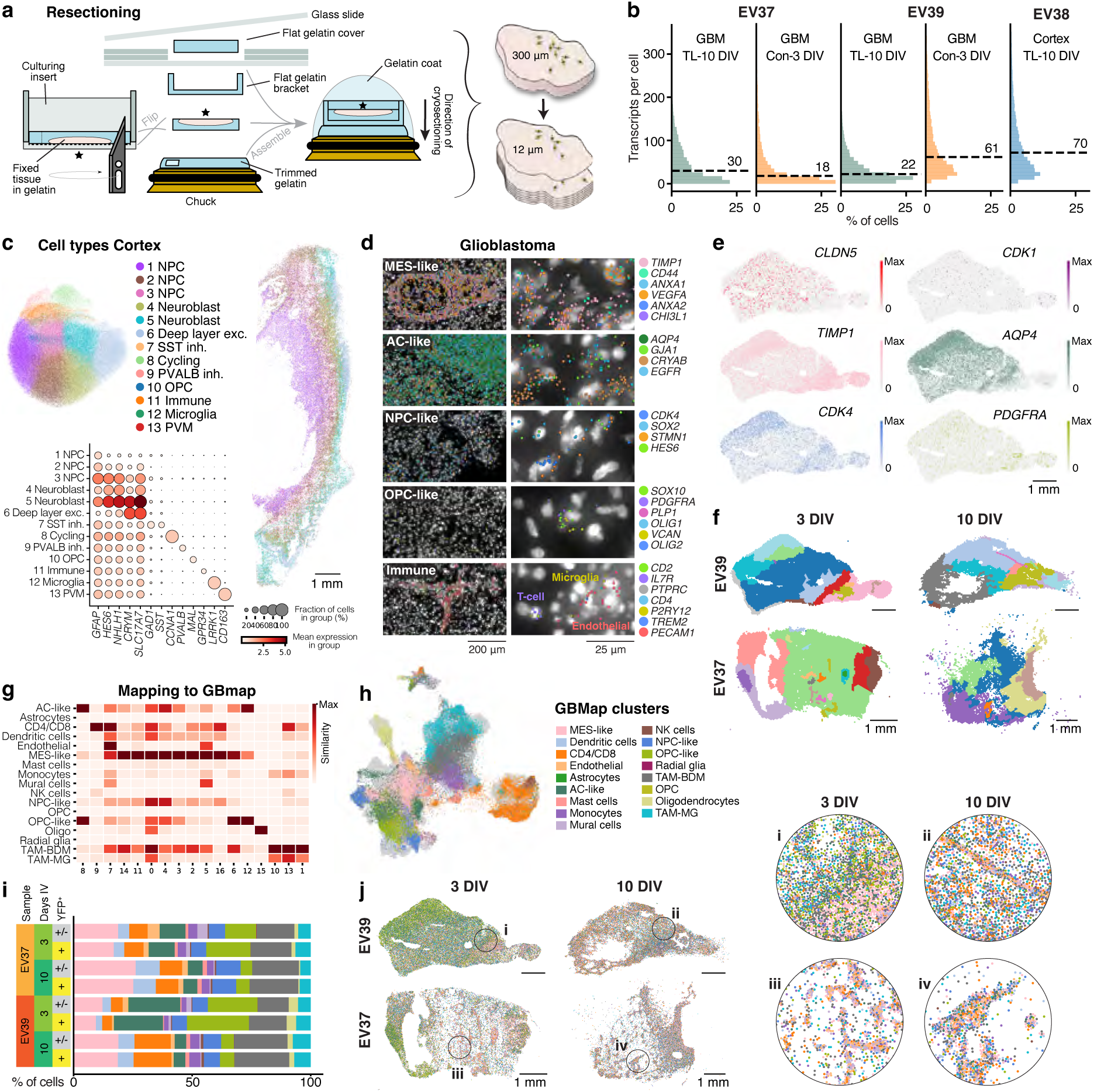
Spatial transcriptomics reveals cellular identities in *ex vivo* cultured samples. **a)** Schematic illustration of our protocol following live timelapse imaging to resection 300 μm tissue slices into 12 μm sections for Xenium spatial transcriptomics. See full protocol in Extended data Fig. 6 and Methods **b)** Distribution of transcripts per cell by sample. IV, *in vitro*. **c)** Cell type annotation of clusters in the cortex based on marker genes. exc., excitatory; inh., inhibitory; NPC, neural progenitor cell; OPC, oligodendrocyte progenitor cell; PVM, perivascular macrophage. **d)** Representative images of gene expression in glioblastoma cell types. Dots represent individual RNA transcripts. AC, astrocyte; MES, mesenchymal; NPC, neural progenitor cell; OPC, oligodendrocyte progenitor cell. **e)** Spatial expression patterns of *CLDN5* (endothelia), *CDK1* (cycling), *TIMP1* (MES-like), *AQP4* (AC-like), *CDK4* (NPC-like) and *PDGFRA* (OPC-like). **f)** Regionalisation of two glioblastoma samples at 3 and 10 days *in vitro*. **g)** Label transfer from GBMap to *ex vivo* glioblastoma samples. **h)** Projection of cells from *ex vivo* glioblastoma to GBMap clusters. **i)** Distribution of cell types between samples, culture time and infection with H2B-EYFP lentivirus. **j)** Spatial distribution of cell types across adjacent glioblastoma slices cultured for 3 and 10 days, from two samples. Colours correspond to the colourkey in h.

**Figure 4:**
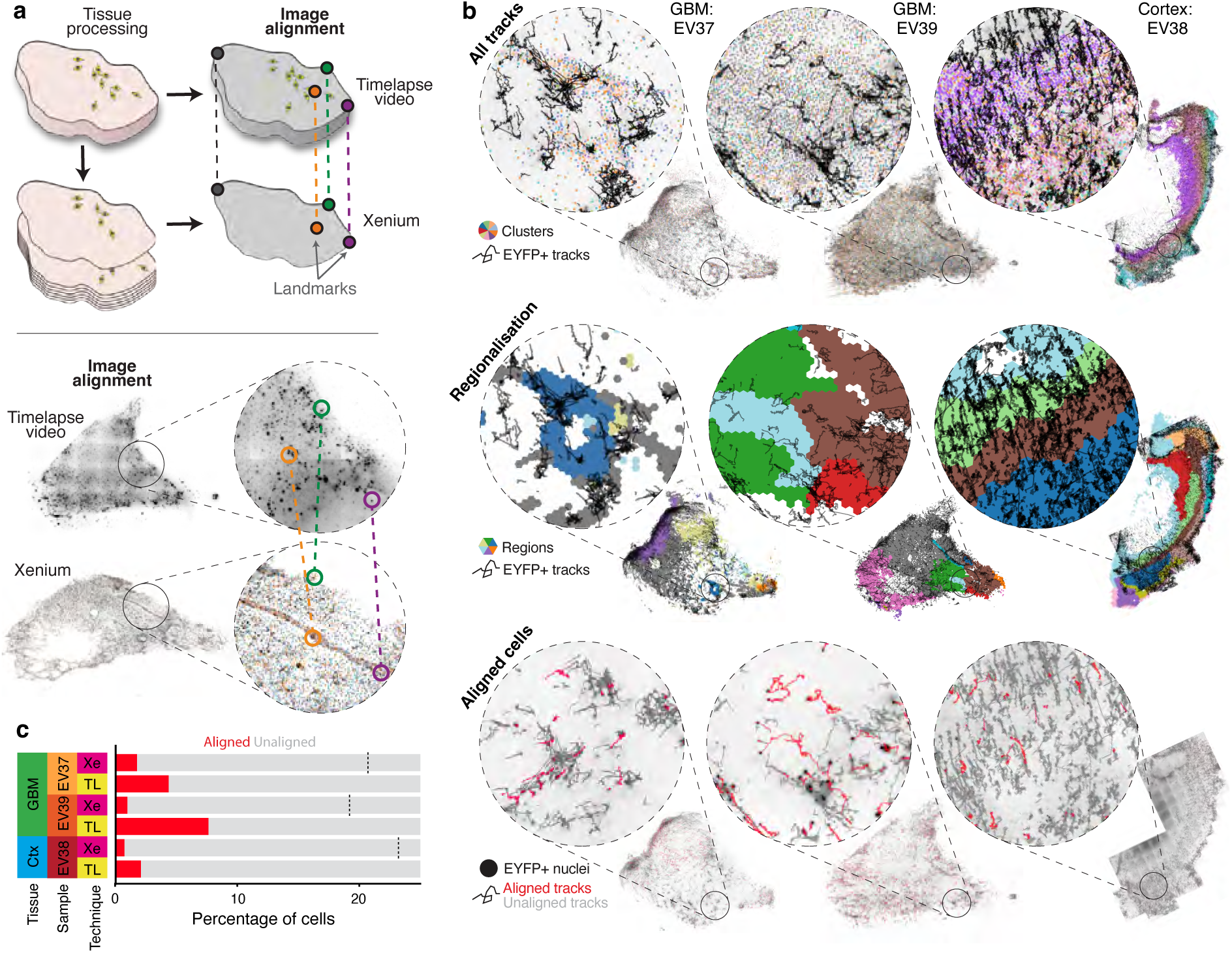
Alignment of *ex vivo* cell tracking to spatial transcriptomics. **a)** After timelapse imaging the thick tissue slices (∼300 μm) are resectioned to 12 μm for spatial transcriptomics. The spatial transcriptomic data is then realigned to the final positions of the timelapse imaging tracks using visual landmarks and a thin plate spline transformation. **b)** Migratory tracks in the context of regional identities. From top to bottom: cellular migration shown overlaid on cluster identities (colours identical to relevant clusters in Figure 3), cellular migration overlaid on regional niches, and cellular tracks of which the expression profile was captured. **c)** The fractions of cells that are aligned from Xenium to live imaging and vice versa. A dashed line indicates the percentage of cells expressing at least one EYFP molecule in Xenium.

We performed spatial transcriptomics of two glioblastoma samples and one fetal cortex following live imaging, at 10 days *in vitro*. We also collected culturing controls from the glioblastoma samples (adjacent vibratome slices) at three days *in vitro*, equivalent to the start of a timelapse experiment. We created a custom 100 gene panel based on known glioblastoma cell type markers, together with the predefined Xenium human brain panel of 266 genes (Table S3).

The tissue quality after live imaging was sufficient for downstream spatial transcriptomics analysis (Fig. 3b), with 70 transcripts per cell detected in fetal cortex and 18 to 61 per cell in glioblastoma. Tissue quality was not systematically different when comparing control samples (3 days *in vitro*) to live imaging samples (10 days *in vitro*). Clustering of the spatial data of the fetal cortex identified stratified layers of neural progenitor cells, neuroblasts, inhibitory- and excitatory neurons, and non-neural populations (Fig. 3c), consistent with our scRNA-seq analysis (Fig. 2b, Extended data Fig. 2).

At three days *in vitro*, glioblastoma explant cultures contained a mixture of immune cells, vascular cells, and cells expressing genes associated with mesenchymal-like (MES-like), astrocyte-like (AC-like), neural progenitor-like (NPC-like), and oligodendrocyte progenitor-like (OPC-like) tumour cells (Fig. 3d-e). Moreover, distinct spatial regions could be identified based on gene expression in tumour sections. This included regions high in MES-like, AC-like, NPC-like and OPC-like associated markers, or highly vascularised regions (Fig. 3e-f).

Next, to annotate the glioblastoma data we performed label transfer from the core GBMap dataset^14^ using scanpy’s ingest function (Fig. 3g-h). Several spatial clusters could be mapped directly to a specific cell type (oligodendrocytes, CD4/8+ T-cells), while many others contained a mixture of transcriptionally related cell types (e.g. tumour cells or tumour-associated macrophages). Clusters of tumour cells were either a mix of AC-like and OPC-like, predominantly MES-like, or a combination of MES-like with other states (Fig. 3g). Interestingly, both AC-like and OPC-like cells showed a decreased abundance after 10 days *in vitro* in favour of a more predominantly MES-like phenotype (Fig 3i) in line with previous observations after *in vitro* culture^15^. Notably, immune cells were able to survive for the duration of the culturing as CD4/8+, natural killer cells (NK), dendritic cells (DC), monocytes, mast cells and tumour-associated macrophages (TAM) were present at both timepoints (Fig. 3i). Furthermore, we observed that the borders between some tumour zones tended to become more diffuse over time in culture (Fig. 3j, i-ii).

Following spatial transcriptomics processing, the samples were stained using DAPI, lectin and a KI67 antibody. These stainings showed a greater number of cycling (KI67+) cells in the cortical sample compared to glioblastoma (Extended data Fig. 7a-c), in line with a higher cell division rate and higher expression of proliferation markers in the fetal cortex (Extended data Fig. 7d-e). Moreover, fetal cortex showed regionalised expression of cycling markers with a clear band extending through the tissue, while cycling cells in glioblastoma were interspersed (Extended data Fig. 7).

### Image alignment between live imaging and spatial transcriptomics

Tissue slices were fixed in PFA immediately after the end of timelapse microscopy to prevent further cell migration between the end of timelapse and start of spatial transcriptomics experiments, which would otherwise hinder image registration. Anatomical features of the tissue, e.g. vasculature and nuclei dense regions, were used to identify landmarks between the timelapse slice and the cryosections, and to perform a thin plate spline transformation using Fiji’s BigWarp^16^ tool (Fig. 4a). This allowed us to map the spatial transcriptomics data and cell identities to the same geometric space of the last timelapse image frame (Fig. 4b), i.e. map migration tracks in relation to cell type clusters and spatial regions. Since the transcriptomic analysis was performed on one to three 12 *μ* m sections, it only contained a subset of cells tracked in the 60 µm imaging depth of the 300 µm tissue slice. In addition, cells that had migrated out of the 60 µm imaging depth were not captured in the collected cryosections. Nonetheless, we were able to map ∼5% of the cells that were tracked during live imaging to a cell in the Xenium data, and up to ∼8% of cells in glioblastoma sample EV39 for which 3 consecutive cryosections were collected (Fig. 4b-c). 1-2% of cells in the Xenium data could be mapped to EYFP+ tracked timelapse cells, due to the sparse labelling used to facilitate cell tracking.

### Results from alignment: cell type-specific behaviours

Cell to cell alignment made it possible to investigate cellular behaviour on a cell type level within a tissue. When comparing aligned to unaligned cells, we found no noticeable differences in cell type composition, indicating that there was no bias in cells aligned (Fig. 5a). We found that most cell types displayed some form of displacement, however this varied by type. For instance, oligodendrocytes in medial areas showed no significant displacement, although there was some concerted displacement with neighbouring cells in oligodendrocytes near the tissue border, presumably because of tissue shrinkage. Conversely, many immune cells (TAM-MG/BDM, CD4/8, monocytes) showed movement throughout the tissue. In the cortex we noticed that most cells remained relatively stationary, while those that did migrate longer distances could be mapped to either immature interneurons or immune cells (Fig. 5b). This is an expected result, given that we imaged the fetal samples with the ventricular zone down, where radial glia and neuroblasts of the excitatory lineages would be expected to move only perpendicular to the imaging plane, whereas interneurons migrate laterally and thus across the imaging plane. The cell and thus across the imaging plane. The cell samples as in the cortex (Fig. 2c, Fig 5c).

**Figure 5:**
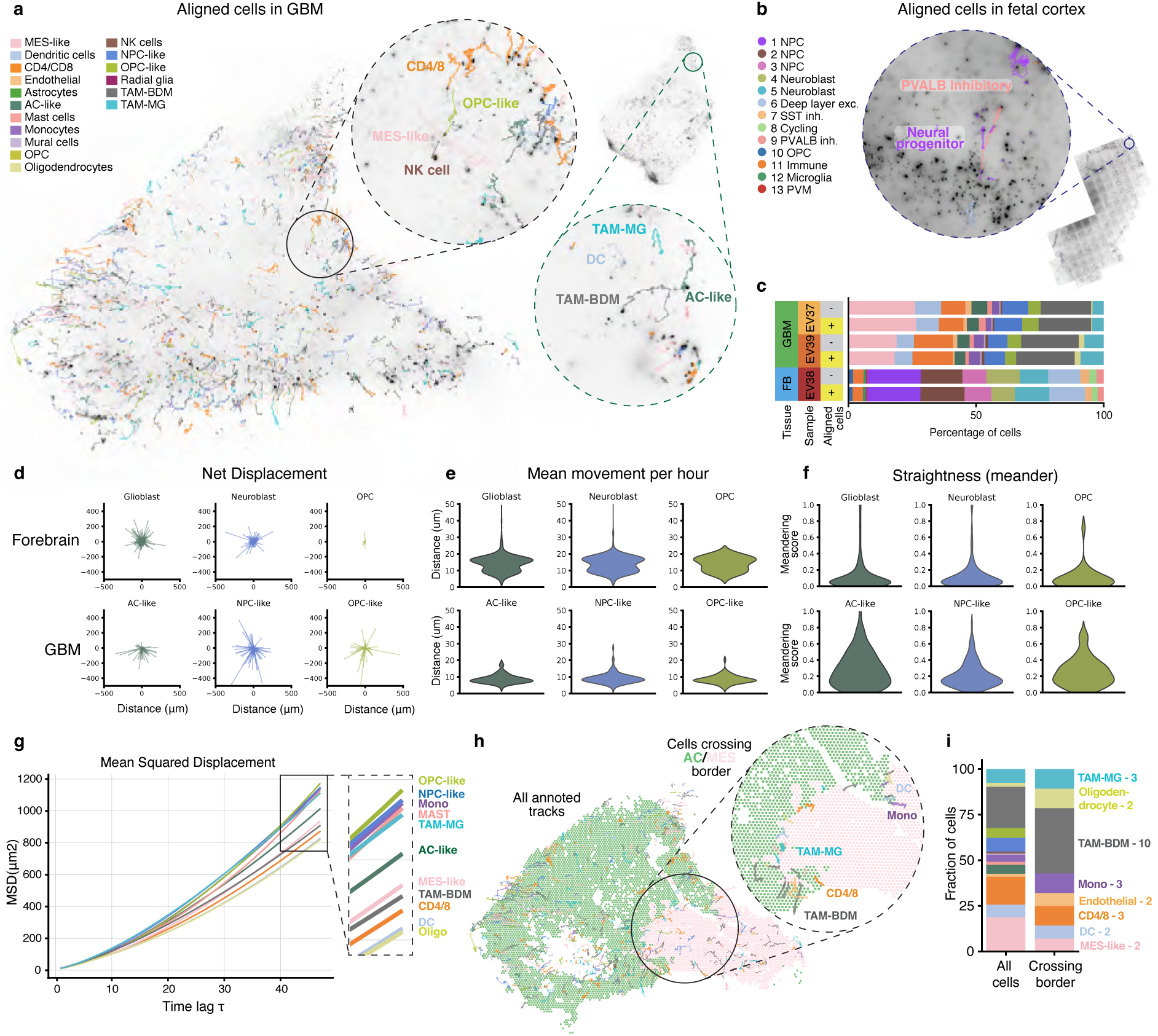
Migration properties of fetal and glioblastoma cell types. **a)** Two glioblastoma samples with cell tracks plotted in the colour of the cell type that they were aligned to in the spatial transcriptomics data. **b)** A fetal cortex sample with cell tracks plotted in the aligned cell type’s colour. **c)** Quantification of cell types of aligned and unaligned cells per sample. **d)** Net displacement of neural progenitors, neuroblasts and OPCs in the fetal cortex and corresponding glioblastoma cell types (AC-like, NPC-like and OPC-like). **e)** Distribution of mean hourly movement per cell for the same cell types. **f)** Distribution of cell meandering scores. **g)** Mean squared displacement for several aligned cell types. **h)** Aligned cell tracks over regions binarized based on a prevalence of AC-like or MES-like expression patterns. The inset shows cell types crossing between tumour regions dominated by AC-like and MES-like gene expression patterns. **i)** Fraction of aligned cells crossing the MES-like/AC-like boundary belonging to each cell type compared to the full population.

We next compared glioblastoma cell states to their closest developmental counterparts in our dataset (AC-like to glioblast, NPC-like to neuroblast, OPC-like to OPC) and found similar behavioural patterns for all groups (Fig. 5d), but with increased displacement and movement in the tumour cells. Notably, while the meandering scores for NPC-like and OPC-like were relatively similar to the fetal cell types, AC-like cells showed a decidedly higher score indicating more targeted movement. Next, mean squared displacement was calculated for each aligned cell type (if there were at least 10 cells). Interestingly, among the tumour cell types, the MES-like subtype showed the lowest overall displacement, while OPC-like and NPC-like cells showed the highest (Fig. 5g). Among the immune cells, monocytes, mast cells and microglia (TAM-MG) showed the largest displacement, while dendritic cells, macrophages (TAM-BDM) and CD4/8 T-cells showed the lowest. Next, we asked to what extent cells cross boundaries between regions of the tumour. We identified a border between a predominantly AC-like and a MES-like region and found that 133 cells crossed this border of which 28 could be annotated (Fig. 5h). 75% (21/28) of the cells crossing this border were immune cells, with TAM-BDM (10/21) being the largest group (Fig. 5g).

### Behaviour around vasculature

Glioblastoma relies on vascular networks to support its rapid growth. For example, perivascular regions provide substrate for cell migration^17^, while hypoxic areas promote a more invasive phenotype^18,19^. In our data, blood vessels were present and preserved throughout the 10-day culturing period, and could be identified based on gene expression, lectin staining, and H&E (Fig. 6a-b, Extended data Fig. 8). Notably, we observed directed migration and an accumulation of cells around vasculature over time (Fig. 6c, Video S2). Most cells in the tumours were located within 500 μm of a blood vessel (Fig. 6d), and cells close to blood vessels tended to migrate along the vessel (Fig. 6e, Video S3). Interestingly, a few cells were also moving perpendicular to the vessel, potentially via diapedesis (Fig. 6e, Video S3-4). Sometimes cells appeared to wait for each other to pass before moving across the vessel, reminiscent of traffic-adapted behaviour (Video S3-4).

**Figure 6:**
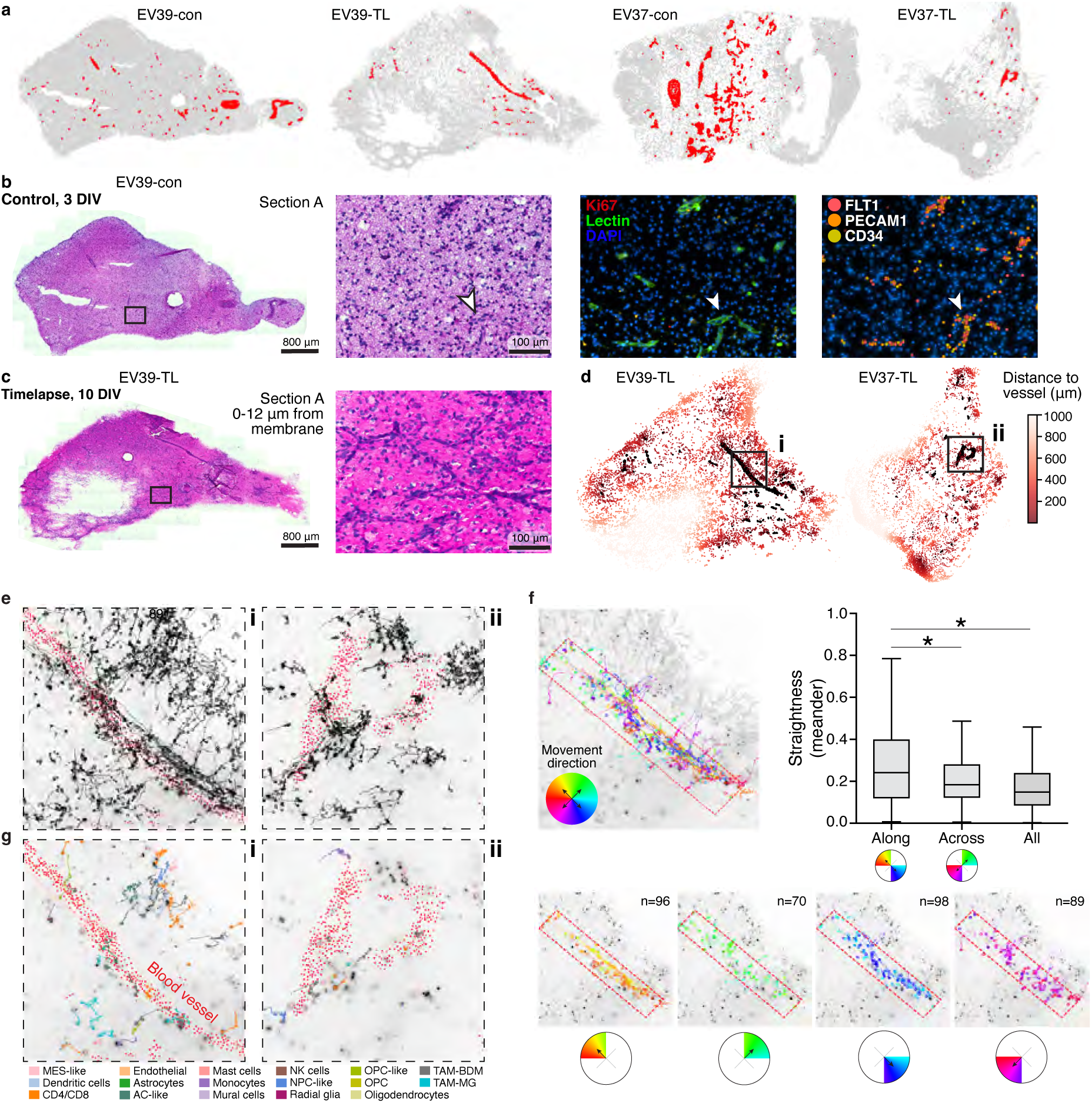
Cell migration behaviour in relation to blood vessels in glioblastoma. **a)** Distribution of blood vessels across the glioblastoma samples at the start (3 days *in vitro*) and end (10 days *in vitro*) of an experiment. con; control, TL; timelapse. **b)** A resectioned 3 days *in vitro* control slice, stained with H&E, Lectin and Ki67, and Xenium spatial transcriptomics. DIV; days *in vitro*. **c)** Adjacent 10 days in vitro slice resectioned post-timelapse stained with H&E. **d)** Distance of cell tracks to the nearest blood vessel at their final position. **e)** Cell tracks around blood vessels. Insets i & ii are from indicated location in D. **f)** All cell tracks intersecting with a blood vessel coloured by the angle of the cell’s net displacement. Tracks are divided into along the vessel (orange/blue) and perpendicular (green/purple). The distribution of the meander score is shown for cells migrating along and across the vessel as well as all other tracks, with cells migrating along the vessel showing a higher score (p<.05). **g)** Aligned cell tracks plotted in colour corresponding to cell type.

To assess the behaviour of cells around vasculature, we asked to what extent cells that migrated along or across the vessel did so in a straight line suggestive of directed migration. Computing the meander index (where a higher index indicates a more straight-line migration), we found that cells migrating along the vessel exhibited a higher meander index, indicating more directed movement as a result of using the vessel as a migratory corridor (Fig. 6f). Conversely, cells migrating across vessels were not different from all cells.

Examining the migration of identified cell types around vessels, we captured a MES-like cell able to migrate both along and across the vessel, an OPC-like cell migrating across, and an AC-like cell migrating along the vessel (Fig. 6g, i). Several immune cells were also migrating in close association with this blood vessel, for example, TAM-MGs, TAM-BDMs, and CD4/8 T-cells (Fig. 6g). We noted a higher expression of vascular and hypoxia-related genes *VEGFA* and *HIF1A* in this large blood vessel (Extended data Fig. 8c), potentially influencing the directed cell migrations observed.

## DISCUSSION

Cellular behaviour is dictated by intrinsic molecular states, perturbed by long- and short-range spatial interactions. Therefore, to understand and explain cell behaviour, methods are needed to simultaneously record dynamic cell phenotypes and the associated molecular states. Arguably, the need for such methods is greatest in human tissues, where our access to *in vivo* experimentation and genetic manipulation is extremely limited.

Ultimately, we would like to record cells in their natural habitat *in vivo* monitoring both cell-cell interactions, cell morphology and complete molecular profiles. While we are far from this challenging goal, much progress has been made. Early methods used physical isolation of single cells. For example, Patch-seq combines temporal recording of cellular electrical responses with cell isolation and single-cell RNA sequencing^20,21^, and has been used to relate electrophysiology, morphology and cell identity in the human brain^22^. Similarly, laser capture microdissection can be applied to cells after timelapse imaging, and used with single-cell RNA sequencing to reveal molecular states and cell identities; this approach was used to discover molecular correlates of cancer drug resistance^23^. One promising emerging technology is Raman microscopy, which noninvasively reveals vibrational spectra of the molecules present in a cell^24^. However, such spectra are difficult to resolve in tissues (compared to 2D culture) and cannot be reliably assigned to identifiable macromolecules such as mRNAs. Conversely, the increased multiplexing of fluorescent reporter proteins (e.g. PICASSO^25^, TMI^26^), live cyclic antibody staining^27^ or *in situ* DNA barcoding^28^ have been used to discover molecular properties of cells after timelapse recording in 2D culture.

However, existing methods are not amenable to use in complex human tissue, even *ex vivo* slice cultures, mostly due to the need to physically access individual cells. Here, we solve that problem by demonstrating how long-term live timelapse imaging can be combined with high-throughput spatial transcriptomics at cellular resolution. The key enabling technical advance was the physical isolation of individual thin cryosections out of the thicker live tissue slice, with preservation of precise cellular alignment. Conceptually, our approach is analogous to how the physical isolation of single cells (e.g. Patch-seq or laser capture microdissection) enabled the use of the full range of single-cell omics methods. Here instead, isolation of single cryosections out of a thicker tissue enables the use of the full range of spatial omics methods. In both cases, linking rich live tissue imaging to rich molecular omics measurements (single-cell or spatial) makes it possible to link cell identity to cell behaviour.

We developed robust protocols for long-term explant culture of both human fetal brain and glioblastoma, and showed that explants remained healthy, continued to proliferate, and maintained rich spatial structure and cell type diversity including immune and vascular cells. We demonstrated that we could track the movement of tens of thousands of cells per sample, for more than one week in culture, and showed that by tracking cells across cell divisions, small lineage trees can be constructed. We subjected cryosections to spatial transcriptomics while maintaining alignment with the timelapse recording, thus linking cell behaviour to cell identity at scale, in human tissue.

OTTR is limited by several methodological constraints. First, accurate lineage reconstruction is limited by the number of mitotic events observed; because human cortical neurogenesis proceeds slowly, longer-term sampling across multiple developmental stages is required to capture deeper branches of the lineage tree. Second, spatial transcriptomics can be performed only after live imaging, precluding dynamic molecular measurements during lineage tracing. Third, resectioning is a challenging manual process, and inherently introduces tissue deformation as a potential source of bias to the alignment procedure. Finally, the inherent heterogeneity of human tissue necessitates large sample sizes to achieve statistical power and reproducibility.

Notably, OTTR is not limited to spatial transcriptomics, but could be used with any spatial omics method that can be applied to a fixed thin tissue section. For example, spatial methods for chromatin accessibility^29^ or modifications^30^, and protein^31^ detection are well established and numerous spatial methods are in development^32^. Long-term explant cultures are also physically accessible and could for example be combined with pharmacological or genetic perturbations, or even optical pooled screens of rich behavioural and molecular phenotypes. Thus, the integration of long-term timelapse recording with spatial omics holds great promise for unlocking some of the mysteries of human tissues.

## AUTHOR CONTRIBUTIONS

Study conception: S.L.; organotypic culturing: E.V., lentiviral infection: E.V.; scRNA sequencing: L.H., E.V.; scRNA data preprocessing: P.L.; scRNA data analysis: E.V., live imaging: E.V., C.M.; microfluidic automation: C.M.; timelapse data processing: C.M.; re-sectioning optimisation: J.J., D.F.G.; re-sectioning for Xenium: J.J., D.F.G., E.V.; immunohistochemistry: I.K.; Xenium optimisation: E.V., J.J., K.T., C.Y.; Xenium data processing: C.M.; data annotation: C.M., E.V.; timelapse and Xenium alignment and analysis: C.M.; data interpretation: C.M., E.V., D.F.G., J.J., S.L.; visualisation: E.V., C.M., D.F.G.; data curation and software: P.L., C.M.; resources: X.H., R.A.B., E.S., X.L., and O.P.; project management: E.V.,; writing - original draft: E.V., C.M., D.F.G., J.J., S.L.; funding acquisition: S.L.

All authors read and approved the final version of the manuscript.

## DECLARATION OF INTERESTS

S.L. is a paid scientific advisor to Moleculent AB, and majority shareholder in EEL Transcriptomics AB (holding patents related to multiplex RNA detection in situ).

## ACKNOWLEDGEMENTS

We thank Stig Linnarsson, Marcus Ridderstolpe and Petroc Klevnäs for manual curation of cell division in timelapse recordings and Razieh Karamzadeh for designing the EYFP reporter lentivirus. We thank Sergio Marco Salas for providing valuable input to the Xenium analysis. We thank the Human Developmental Biology Resource (www.hdbr.org) for access to fetal tissues. We acknowledge support from the National Genomics Infrastructure in Stockholm funded by Science for Life Laboratory, the Knut and Alice Wallenberg Foundation and the Swedish Research Council, and NAISS for assistance with massively parallel sequencing and access to the UPPMAX computational infrastructure. We acknowledge the In Situ Sequencing Facility at SciLifeLab, funded by Science for Life Laboratory and the Swedish Research Council, for providing in situ sequencing services.

We thank Wei Li for aiding in the photographic documentation of the sample embedding protocol. This work was supported by grant from Erling-Persson foundation (Atlas of Childhood Disease), Torsten Söderberg foundation, Knut and Alice Wallenberg foundation (2023.0340, 2018.0172, 2018.0220), Swedish Research Council (2022-01248) to S.L.. The work done in Cambridge was supported by the National Institute for Health and Care Research (NIHR) Cambridge Biomedical Research Centre (NIHR203312). The views expressed are those of the authors and not necessarily those of the NIHR or the Department of Health and Social Care.

## SUPPLEMENTARY VIDEO LEGENDS

**Video S1: Cell divisions.** Representative field of view in a glioblastoma tissue section with cell divisions indicated by red arrows.

**Video S2: Cell accumulation around vessels**. Red arrow highlights a blood vessel where cells accumulate during culturing of glioblastoma sample EV39.

**Video S3: Cell migration around a large blood vessel.** Area of sample EV39 corresponding to Figure 6. Migration is observed along a large blood vessel as well as perpendicular to it.

**Video S4: Zoom in of cells crossing and following a blood vessel.** Two examples of cellular migration in relation to a blood vessel are shown: a) a cell entering the blood vessel and following it for its migration and b) a cell repeatedly crossing the vessel.

## METHODS

### STUDY PARTICIPANT DETAILS

#### Donors

Human prenatal samples were collected from elective medical abortions at the Department of Gynecology, Danderyd Hospital and Karolinska Huddinge Hospital, Addenbrooke’s Hospital in Cambridge, and the Human Developmental Brain Resource following oral and written informed consent by the patient. In Sweden, the use of abortion material was approved by the Swedish Ethical Review Authority and the National Board of Health and Welfare (2018/769-31 with amendments). In the UK, approval was obtained from the National Research Ethics Committee East of England, Cambridge Central, and from the North East – Newcastle & North Tyneside 1 Research Ethics Committee (Local Research Ethics Committee, 96/085; DNR2019-04595).

#### Patient samples

Human glioblastoma samples were collected from the Karolinska Hospital with informed consent from the patients and with ethical approval from the Swedish Ethical Review Authority (2020-03505). The use of samples was approved by the Swedish Ethical Review Authority (2020-02096).

Details about all samples can be found in Table S1.

## METHOD DETAILS

### Tissue sample collection

#### Human fetal forebrain

12 samples of human prenatal forebrain were used in this study, at post-conception weeks (PCW) 7-10. The post-conception age of the fetuses was estimated by information from the clinical ultrasound, last menstrual period, true crown-rump-length, and age-dependent anatomical landmarks.

For samples collected at the Karolinska Hospital, the tissue was immediately transported to the laboratory following abortion and dissected in ice-cold 0.9% NaCl solution under sterile conditions within 1-2 hours post abortion. The forebrain was dissected and kept in ice-cold Hibernate-E medium (ThermoFisher #A1247601) until further processing.

For samples collected in Cambridge, tissues were dissected in a class II hood on the day of collection and stored in Hibernate-E medium at 4 °C. The tissue was shipped to Sweden at refrigerated temperature and delivered two days after abortion. The procedure is covered under ethics REC: 96/085.

#### Human glioblastoma

Nine glioblastoma samples were collected from patients. During surgery, the glioblastoma was placed in Hibernate-E medium on ice and transported as such. Upon arrival, samples were washed in fresh Hibernate-E medium and kept in ice-cold Hibernate-E medium until further processing.

### Organotypic slices

#### Primary glioblastoma tumours

The full detailed protocol is available at https://dx.doi.org/10.17504/protocols.io.8epv5op9jg1b/v1, including appropriate safety measures. Briefly, glioblastoma samples were cut into well-shaped chunks or strips of tissue and cleaned up from any necrotic areas and loose blood vessels, while kept in ice cold carbonated tissue* (see “Media compositions” below). Put 5-7 mL of 2% low gelling agarose (or 1%/3% if the tumour sample is very soft/hard) (Thermo Scientific #J66319.14 or ThermoFisher #16520050) into a plastic embedding mould and chill on ice for 1 min. Place 2-3 strips of tissue vertically into the gel and swirl gently a couple of times. Leave on ice for another 2-5 minutes until the agarose has hardened. Put sticky tape on the vibratome platform and glue the agarose chunk with tissue on the tape. Put the platform inside the vibratome chamber and fill it with carbonated tissue processing medium until it covers the agarose. Section at 300 μm. Usually a speed of 0.2 mm/s was used but adjusted according to the stiffness of individual glioblastoma tumours (more stiff meaning slower sectioning speed). Use a microspatula or glass suction tube to collect individual slices into a 12-well plate on ice, containing tissue processing medium. Change tissue processing medium once while sectioning.

#### Fetal forebrain

Dissected forebrains were processed around 6 to 48 hours after tissue collection, depending on the source (Karolinska Hospital/the UK). Tissues were stored at 4 °C in Hibernate-E medium during transport from the UK and until processing. Upon arrival, Hibernate-E medium was replenished, and forebrains were dissected in Hibernate-E medium in a Petri dish on ice. Forebrains were cut into strips along the dorso-ventral axis and cut at the intersection between the (thin) cortex and (thick) ganglionic eminences.

### Lentiviral infection and culturing

Heat slice maintenance medium^#^ in a 37°C water bath. Meanwhile, wash the forebrain/glioblastoma slices twice for 10 min while gently rocking at room temperature, once in tissue processing medium* then in slice washing medium^%^.

Transfer vials of lentivirus from a −80°C freezer on dry ice, and thaw on ice before usage. We used a plasmid with H2B-EYFP under a hPKG promoter causing EYFP expression and fluorescence in cell nuclei. The plasmid was packaged into a VSVG generation 3 lentivirus, ultrapurified and at a titre of 10^9^ (Vectorbuilder), causing ubiquitous infection. A H2B-GFP construct was previously also tried and worked fine but with slightly lower fluorescence intensity in nuclei. For a virus titre of 10^9^, dilute 2 μL (for glioblastoma slices) or a range of 2-3.5 μL (cortical slices) in slice maintenance medium to a final volume of 500 μL per well, in a 12-well plate. Remove the tissue processing medium from the slices and add the virus-medium mix, one well at the time. Put the plate in a 5% oxygen (hypoxic) incubator overnight. The next morning, pre-heat slice washing medium and slice maintenance medium. Remove the virus-containing medium from the slices and do a quick rinse with 1 mL slice washing medium per well. Wash with 1 mL slice washing medium twice for 10 min while rocking at room temperature. Then wash with 1 mL of 1:10 diluted slice maintenance medium in MEM, for 10 min while rocking. Remove and add the same medium again.

While the slices are washing, prepare 6-well plates for culturing. Add 1 mL of slice medium to a well and lower a PTFE-membrane insert using sterile forceps. Angle the plate towards you to ensure any bubbles rise to the side of the insert. Carefully put tissue slices in the middle of the PTFE-insert and remove excess liquid around the edges of the slice. Change maintenance medium the day after plating, then every 48h. Leave plate for 48-72 h in the incubator (or until fluorescent signal from the lentivirus is apparent) before imaging.

#### Media compositions

Media compositions:
| **Tissue processing medium\*** |  |  |  |
| --- | --- | --- | --- |
| **Media component** | **Volume for 250 mL** | **Company** | **Cat. number** |
| DMEM (high glucose, no glut) | 242.5 mL | ThermoFisher | #11960044 |
| HEPES 1M | 2.5 mL | ThermoFisher | #15630080 |
| Anti-Anti 100X | 2.5 mL | ThermoFisher | #15240062 |
| GlutaMAX 100X | 2.5 mL | ThermoFisher | #35050061 |
| **Slice maintenance medium<sup>#</sup>** |  |  |  |
| **Media component** | **Volume for 20 mL** | **Company** | **Cat. number** |
| MEM (no glutamine, phenol red*) | 12.54 mL | ThermoFisher | #51200038 |
| HBSS | 5 mL | ThermoFisher | #14025092 |
| 33% D-Glucose | 260 µL | Merck | #G8769 |
| HEPES 1M | 200 µL | ThermoFisher | #15630080 |
| Human serum (heat inactivated, sterile filtered) | 1 mL | Merck | #H5667 |
| B27 supplement 50X | 400 µL | ThermoFisher | #A3582801 |
| Anti-Anti 100X | 200 µL | ThermoFisher | #15240062 |
| N2 supplement 100X | 200 µL | ThermoFisher | #17502048 |
| GlutaMAX 100X | 200 µL | ThermoFisher | #35050061 |
| **Slice washing medium<sup>%</sup>** |  |  |  |
| **Media component** | **Volume for 20 mL** | **Company** | **Cat. number** |
| Anti-Anti 100X | 200 µL | ThermoFisher | #15240062 |
| GlutaMAX 100X | 200 µL | ThermoFisher | #35050061 |
| MEM (no glutamine, phenol red) | 19.6 mL | ThermoFisher | #51200038 |

\*We used MEM without phenol-red as it interferes with optical clarity for imaging.

### Cell dissociation

Ice cold Earle’s Balanced Salt Solution (Worthington) was carbogenated (95% O2/ 5% CO2) and used throughout the whole procedure. Cortex and glioblastoma slices were dissociated using the Worthington’s Papain Dissociation System (Worthington)(Protocols.io: https://dx.doi.org/10.17504/protocols.io.xmbfk2n). Tissues were enzymatically digested at 37 °C for 10-15 min, followed by trituration using fire polished glass Pasteur pipettes. Cell suspensions were filtered through a 30 μm cell strainer (CellTrics, Sysmex) and centrifuged for 5 min at 200 g to obtain cell pellets. Supernatants were carefully removed, and cells resuspended in small volumes of EBSS (depending on cell density). Cell concentrations were estimated using a counting haemocytometer (Bürker/Neubauer chamber) and diluted with EBSS until the desired concentrations were reached. All suspensions were kept on ice until loading on the 10X Chromium chips.

### Single-cell RNA sequencing

Droplet-based single-cell RNA sequencing was performed using the 10X Genomics Chromium Single Cell Kit v3. Single-cell suspensions concentrated at 800-1200 cells/μL were mixed with master mix and nuclease free water according to the Chromium manual, targeting 5000-10000 cells per reaction. 11-12 PCR cycles were used for cDNA synthesis, and the rest of the library preparation was performed according to the manufacturer’s instructions (10X Genomics, Illumina). All libraries were sequenced on an Illumina NovaSeq 6000 using S4 to a target sequencing depth of 100,000 reads/cell. Sequencing saturation was examined for each sample using preseq (https://github.com/smithlabcode/preseq).

### Stagetop incubation

Tissue was kept alive for prolonged time periods using a combination of temperature maintenance, gas exchange and continuous replacement of culture media. Tissue was kept in an Okolabs electrically heated chamber for Nikon Motorized Stages (H301-NIKON-TI-S-ER) with a consistent internal media temperature of 37 ℃ as measured with a temperature probe inserted and held in place using a 3D printed holder at the bottom of a 1X PBS filled well adjacent to the sample well. The chamber had a Koehler lid (H301-KOEHLER-LID) at the top to allow brightfield illumination and was closed at the bottom with a heated metal holder for 6-well plates (providing open space for imaging). Connected to the chamber was a premixed gas source (5% O2, 5% CO2, Strandmöllen) that was limited to an output of 0.2 bar and humidified using the Okolabs gas controller (UNO-T-H-PREMIXED). An Okolabs chamber riser (MICROFLUIDIC-C-RISER) was added to the stagetop chamber with 2×6 insertion holes for 5/32 inch (∼4.2 mm) external diameter tubing (closable by M5 grub screw). For each cultured sample two holes were opened (one on each side of the chamber) to route inlet and outlet tubing (1.6mm ID, 3.2mm OD; Tygon3350, VWR228-4331) to blunt insertion needles in a 3D printed insert holder to both deposit and remove media. The outlet needle was bent to be placed at equal height to the insert membrane to maintain a consistent level of media. To prevent backflow from the outlet back into the chamber, a polysulfone check valve (West Group; PSCV E-C-180017) was installed at the chamber exit. Tubing was connected to a Tecan Cavro XCalibur syringe pump with a 9 port ceramic distributor head (XC9; 20737494) using a glass 5 mL syringe (20725593). The pump’s ceramic distributor head connected the inlets/outlets of the tissue chamber with the culture media and waste bottle. Every 4 hours the pump would first distribute 100 mL of fresh media to the chamber, then remove any excess media above the tissue line, after this the ceramic head was automatically washed with 70% ethanol for 5 minutes before being rinsed with deionized water. This ensured that no growth would take place in the pump head, nor would any remaining ethanol be distributed to the tissue.

### Live timelapse imaging

All imaging was conducted on a Nikon TiE2-Eclipse using an ANDOR Sona 4.2B-11 large FoV (25 mm) camera using a Lumencor SPECTRA III (VBCTGYRnIR; 90-10323) light source. For overview images we used an CFI S Plan Fluor ELWD 20X objective (N.A. 0.45; MRH08230) while for the live imaging we used a CFI S Plan Fluor ELWD 40X objective (N.A. 0.6; MRH08430). When imaging EYFP fluorescence, excitation and emission filtering was done using a Semrock LED-CFP/YFP/mCherry-3X-A filter cube (32mm; MXR00756). Prior to live imaging each slice was screened under the microscope for the presence of EYFP^+^ cells. A standardized NIKON job was used to set up live imaging. First, a brightfield overview was taken for each section and a region of interest (ROI) selected based on this image, ideally comprising the full tissue slice. Next, for each slice the Z-coordinates of tissue-membrane interface was determined at each corner of the ROI and a Z-stack was set for imaging going from −15 to +60 μm of the lowest point. Next, a Z-stack was taken in a separate well filled with culture media to generate a background image used for correction of vignetting in image corners and limited fluorescence from the media. The imaging loop is then started by autofocusing on the first position using a reduced exposure time (4% laser strength, 60ms, 4×4 binning) to adjust the z-stack for changes in the media volume. Each position is imaged at 25% laser strength with 500 ms exposure time. a new iteration of the imaging loop is started at the 15 minute mark. At the end of each day all the collected images are uploaded to the computing cluster without disrupting the ongoing job. After termination of the job a brightfield overview image of the tissue is taken at 20X magnification and the tissue is immediately fixed for 30 minutes with 4% PFA. Each iteration of the imaging loop is saved as a separate file and processed using a custom python script. First, images are read into python using nd2reader (https://github.com/Open-Science-Tools/nd2reader) and for each Z-stack the maximum projection is calculated, and vignetting is corrected for by dividing the image by the collected background tile. Values are capped at the 1000^th^ quantile. Images are then stitched with a 10% overlap between positions and saved as tiff files. For visualization purposes videos of each position and stitched tissues are exported in .mov format, where stitched images are binned 2×2 if they consist of 17 or more tiles and 4×4 if they consist of 65 or more (these videos are not used for analysis).

### Resectioning for spatial omics

To ensure that the cells tracked during live imaging are captured in spatial transcriptomics data and that the corresponding images can be aligned, spatial transcriptomics data must be obtained from the bottom-most, PTFE facing, regions of the slice culture without significant tissue warping. To account for this, we have tested: staining and imaging the slice culture without removing it from the insert membrane, either with tissue clearing (MERFISH protocol with prolonged clearing and additional permeabilization using SDS and Triton X-100) or without (RNAscope, osmFISH protocol with added branched DNA amplification), removing the slice culture from the insert membrane (mechanically or enzymatically), either for free-floating staining or for capture on glass surface (same smFISH protocols as above), and found these approaches to be unsuitable due to poor staining or image quality in the region of interest (epifluorescence and confocal microscopes were tested), damage to the tissue or warping, unreasonably low throughput, or a combination of these factors. Therefore, we have chosen to cryosection the slice cultures into thin slices more compatible with conventional smFISH protocols. To optimize this step, we have tested: the need for tissue fixation (with different fixation times) and cryopreservation, tissue quality at different culturing timepoints, different embedding media (gelatin and sucrose solutions at varying concentrations, O.C.T., and mixtures thereof), and corresponding cryosectioning temperatures. The protocol outlined below yielded most consistent and well aligned tissue sections from the immediate membrane-facing region of the slice culture.

Samples were briefly washed with 1× PBS (ThermoFisher #AM9625) and fixed with 4% formaldehyde (ThermoFisher #28908) in 1× PBS for 15 min. Solutions were added both underneath and atop the insert membrane, but caution must be taken not to disrupt the tissue. After fixation, the tissue was washed five times with 1× PBS for at least five minutes each. Samples were cryoprotected by washing with 15% sucrose (Sigma Aldrich #84097) in 1× PBS for two hours, followed by 30% sucrose in 1× PBS overnight at 4 °C. Samples that had been used for timelapse imaging were embedded in a solution of 12% porcine skin gelatin (Sigma Aldrich #G1890) in 1× PBS with 30% sucrose (pre-warmed to 37 °C), positioning the tissue for cutting parallel to the membrane surface, and frozen on dry ice (Extended data Fig. 6; for full protocol see https://dx.doi.org/10.17504/protocols.io.8epv5op9jg1b/v1). Note that gelatin embedded tissue samples require lower cryosectioning temperatures and pre-warmed capture slides. Samples which had not been used for timelapse imaging and therefore did not require high-precision positioning were embedded in O.C.T. (Sakura #4583) in custom-made embedding moulds that fit within the culture insert and frozen on dry ice.

### Xenium *In Situ* Gene Expression

Fixed frozen 12 µm tissue sections were placed on pre-warmed Xenium capture slides. The tissue was fixed and permeabilized according to the Xenium Fixation and Permeabilization Protocol (Demonstrated Protocol CG000581) with the following adaptation: the fixation time was reduced to 10 minutes. A pre-designed Human Brain panel (266 genes) and a custom gene panel (100 genes, Table S2) were applied to the tissue. Probes were hybridized to the target RNA, followed by ligation and enzymatic amplification to generate multiple copies of each RNA target, as outlined in the Probe Hybridization, Ligation, and Amplification User Guide (User Guide CG000582). The prepared Xenium slides were then loaded onto the Xenium Analyzer for imaging and analysis, following the Decoding and Imaging User Guide (User Guide CG000584). Instrument software version 1.6.1.0 and software analysis version 1.6.0.8 were used.

Xenium Slides & Sample Prep Reagents, #1000460

Xenium Decoding Reagents, #1000461

Xenium Decoding Consumables, #1000487

Xenium Human Brain Gene Expression Panel, #1000599

## QUANTIFICATION AND STATISTICAL ANALYSIS

### scRNA-seq data preprocessing

Illumina runs were demultiplexed with cellranger mkfastq version 6.1.2 (10x Genomics). Read mapping and unique molecular identifier (UMI) counts were determined using STARSolo^33^ version 2.7.10a, using human genome GRCh38.p12 and transcript annotations from ENSEMBL release 93 from reference package GRCh38-3.0.0 as available from 10Xgenomics (www.10xgenomics.com). STARSolo was run with the following parameters:

--soloType CB_UMI_Simple

--soloCellFilter EmptyDrops_CR <EXPECTEDNCELLS> 0.99 10 45000 90000 500 0.01 20000 0.01 10000 --soloCBmatchWLtype 1MM_multi_Nbase_pseudocounts

--soloUMIfiltering MultiGeneUMI_CR

--soloUMIdedup 1MM_CR

--clipAdapterType CellRanger4

--outFilterScoreMin 30

Replicates were pooled, resulting in one loom file (loompy.org) per sample.

### Quality control

Samples were analysed with “cytograph qc” (https://github.com/linnarsson-lab/adult-human-brain), which uses a modified version of DoubletFinder to calculate a doublet score for each cell^34^. Cells were filtered based on their total number of mRNA molecules—as counted by unique molecular identifiers (UMIs)—and percentages of unspliced RNA, as well as doublet scores. Cytograph qc was run with standard parameters, where cells with fewer than 1000 UMIs, unspliced molecule fraction less than 0.1, or a doublet score below 0.4 were removed from further analysis.

### Clustering

Cells were clustered using an updated version of the Cytograph package (https://github.com/linnarsson-lab/adult-human-brain). All cells that passed QC for samples XDD:394, XDD:398, and XDD:415 (Table S1) were pooled into a single dataset for clustering. The command “cytograph build” was run with configuration params: batch keys: Donor, features: variance, steps: nn, embeddings, clustering, aggregate, export. In this case batch correction was done per tissue donor. All cells that passed QC for XDD:420 were pooled into a single dataset for clustering. The command “cytograph build” was run with configuration params: batch keys: SampleID, features: variance, steps: nn, embeddings, clustering, aggregate, export. In this case batch correction was done per experiment (4 vs 10 days *in vitro*). Default values were used for other parameters.

### Annotation

Clusters were manually annotated based on literature, known markers of neurodevelopmental cell types. We computed cell cycle scores as previously described^35^, using the expression of a set of well-known cell cycle genes as a proxy for active proliferation and calculating the cell cycle score as the fraction of UMIs those genes represented, and then the percentage of cells per cluster with a cell cycle score > 0.01.

Code available in deposited Jupyter Notebooks (https://github.com/linnarsson-lab/OTTR).

### Cell tracking and calculation of track measures

Cell segmentation was conducted using an adapted watershed segmentation method.. Cells were tracked using btrack, a bayesian cell tracking algorithm^11^ with the following settings for the motion model: Probability unassigned - 0.001; maximum frames lost - 5; P_sigma - 15; G_sigma - 5; R_sigma - 10; and for the hypothesis model: lamda_time - 5; lambda_dist - 3; lambda_link - 10; lambda_branch - 50; theta_dist - 30; dist_thresh - 30, apoptosis_rate - 0.001 and using the ‘relax constraints’ settings for the optimization problem. The algorithm contains an additional step to correct for local background fluorescence. Next, all the generated tracks were exported to a Shoji workspace^36^ as a rank-3 tensor of ‘cells’ by ‘coordinates’ by ‘time’. Next, for each track we calculated several distance measures including total distance travelled, net distance travelled (distance between start and end point), maximum distance from start position, meandering index (net distance divided by total distance), eccentricity of the nucleus at every position, movement inconsistency (mean residual from mean displacement) and mean squared displacement per track. The mean squared displacement could then be used to estimate the diffusion coefficient of a tissue, which in turn was used to model diffusion of cells over longer time periods simulating diffusion in patients.

### Processing of Xenium’s spatial datasets

Xenium experiments were processed using Xenium Ranger (version 1.6.0.8) with default settings. Next, Scanpy (version 1.9.3) was used to Cluster and annotate the data. For the Glioblastoma samples we filtered out cells with less than 15 counts or 5 unique genes. Data was log-scaled, 10 principal components were as input for the nearest neighbour graph based on the 15 nearest neighbours. A resolution of 1 was used for the louvain clustering. For the cortical sample a cut-off of 100 counts and 40 genes was used, with 50 principal components and a cluster resolution of 0.6. Clusters were annotated based on known markers for neurodevelopmental cell types including *HES6, NHLH1* (neuroblast), *SLC17A7* (excitatory), *GAD1* (inhibitory), *SST, PVALB* (interneuron), *CCNA1* (cycling marker), *MAL* (oligodendrocyte precursor cells), *LRRK1* (microglia), *CD163* (perivascular macrophages). Conversely, glioblastoma samples were annotated by transferring labels from the GBMap core scRNA-seq dataset^14^ using scanpy’s ingest function, which uses a KNN classifier trained on the reference dataset to predict labels. To train the classifier we restricted both datasets to overlapping genes. Finally, regionalization in Xenium sections was identified using FISHscale^37^ both on the untransformed and transformed unfiltered Xenium transcripts.

### Alignment of Xenium and timelapse microscopy

The Xenium data was aligned to the time lapse microscopy in two ways. First, using anatomical landmarks, the H&E stained Xenium section was morphed into the space of the last timelapse imaging frame, allowing pixel-level transformations between the two coordinate systems. Second, EYFP-positive cells in the Xenium section were matched with tracked EYFP-positive cells in the last timelapse imaging frame. In more detail, pixel-level morphing was done using Fiji’s BigWarp function^16^. We used the final timestep of the live imaging series as the target image and the H&E-stained section used for Xenium as the moving image. We then identified roughly 30 landmark positions across the two images based on tissue morphology, e.g. blood vessels or dense areas, and the tissue’s borders and used these to anchor the thin plate spline transformation. This transformation was then used to transform both the cell centroids and the RNA coordinates of the associated Xenium experiment. Next, given deformations of the tissue due to processing we did not directly link EYFP fluorescent cells from the timelapse imaging to the nearest Xenium cell. Instead, we first overlapped the H&E with an image of EYFP fluorescence in the tissue after the xenium processing taken on the Nikon TiE2 (also aligned using BigWarp). We used watershed segmentation to identify EYFP fluorescent cells and filtered out all other cells from cell-to-cell alignment. For cell-to-cell alignment we identified each cell tracked in the time lapse imaging present at the final imaging step and identified its nearest EYFP-positive neighbour in the Xenium section with a maximum distance of 50 μm. Following label transfer of the Xenium datasets to timelapse datasets, tracking measures were stratified by cell type and distance to nearest blood vessel was calculated by first defining blood vessels as consisting of any endothelial cell or pericyte that has at least 4/10 nearest neighbours of these groups and then calculating the minimum distance from any cell to any vascular cell.

## DATA AVAILABILITY

- scRNA-seq BAM files are available from the European Genome/Phenome Archive (https://ega-archive.org/) under accession number TBA.
- Xenium *in situ* data and images are available from the BioImage Archive (https://www.ebi.ac.uk/bioimage-archive/) under accession number TBA
- scRNA-seq count matrices and Xenium *in situ* data are also available from our companion GitHub page at https://github.com/linnarsson-lab/OTTR.
- Any additional information required to reanalyse the data reported in this paper is available from the corresponding author upon request.

## CODE AVAILABILITY

- All original code for the analysis and visualisation of data has been deposited to our companion GitHub page at https://github.com/linnarsson-lab/OTTR.

## Notes

https://github.com/linnarsson-lab/OTTR

