## Supplemental Figures for "Organotypic Timelapse recording with Transcriptomic Readout (OTTR) links cell behaviour to cell identity in human tissues"

**This PDF contains:**

**Extended Data Figures 1-8**
**Extended Data Figures 1-8 figure legends**

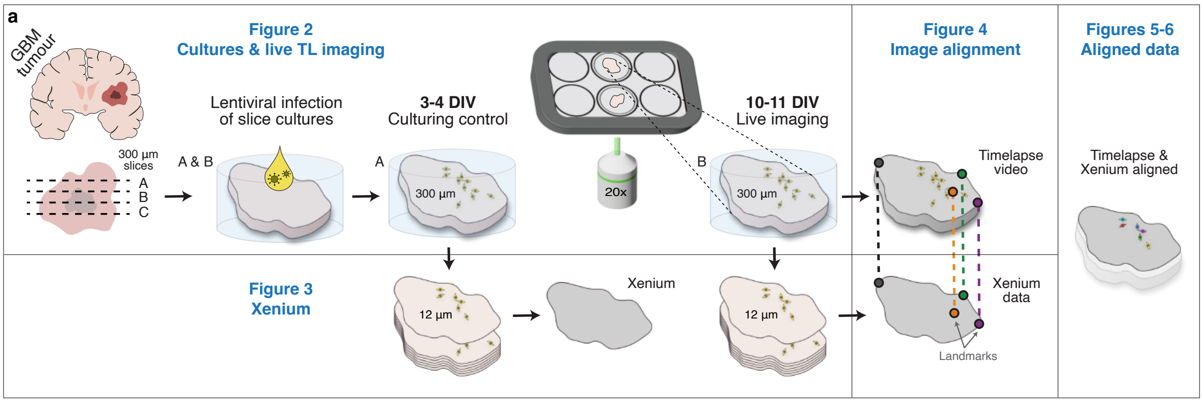


**Extended Data Fig. 1: Schematic overview of the method and associated Figures. a)** Schematic illustration of the whole workflow. Boxes indicate which part of the method is presented in which Figures.

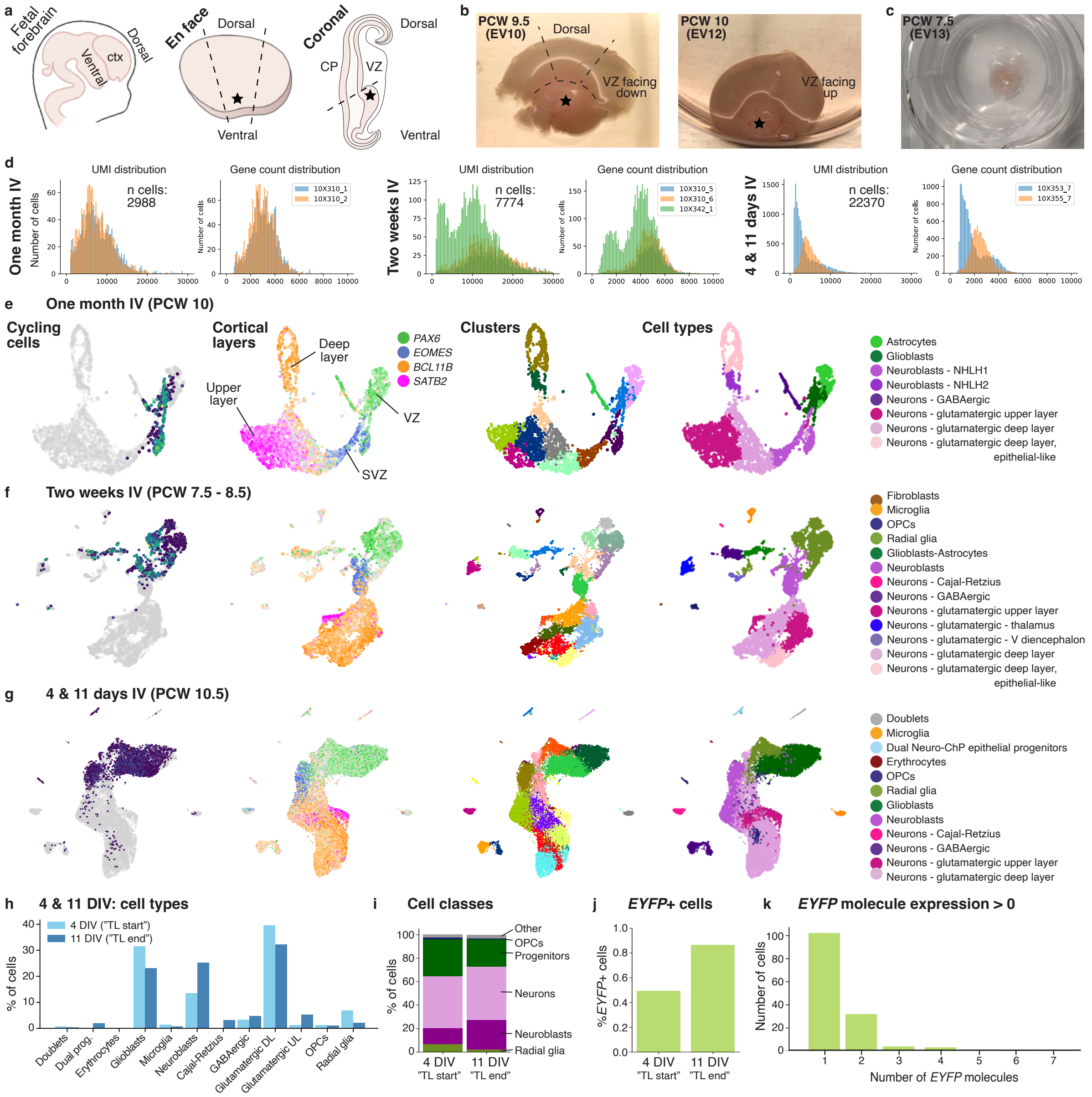

**Extended Data Fig. 2: scRNA-seq validation of cultured fetal cortex explants. a)** Schematic of fetal forebrain anatomy. Ctx; cortex. Dashed lines indicate how they were sectioned, and stars point to the ventral forebrain. **b)** Images of two forebrain samples, showing one hemisphere. **c)** Sample placed ventricular zone side down on a PTFE membrane of a culturing insert. Imaging was always performed on the cortical parts. **d)** UMI and gene count distribution from scRNA-seq data of samples cultured for one month, two weeks, and four and 11 days *in vitro*. IV; *in vitro*. **e)** scRNA-seq UMAPs of cortex explant cultured for one month, coloured by (left to right) cell cycle score, markers for cortical layers, clusters, and annotated cell types. **f)** Same as B) but two weeks *in vitro*. **g)** Same as B) but four and 11 days *in vitro*. **h)** Percentage of cell types at four and 11 days *in vitro*, the equivalent of when a live timelapse imaging experiment started and ended. TL; timelapse. **i)** Percentage of cell classes at four and 11 days *in vitro*. **i)** Percentage of *EYFP*-positive cells at four and 11 days *in vitro*. **k)** Number of *EYFP* molecules in cells positive for *EYFP*.

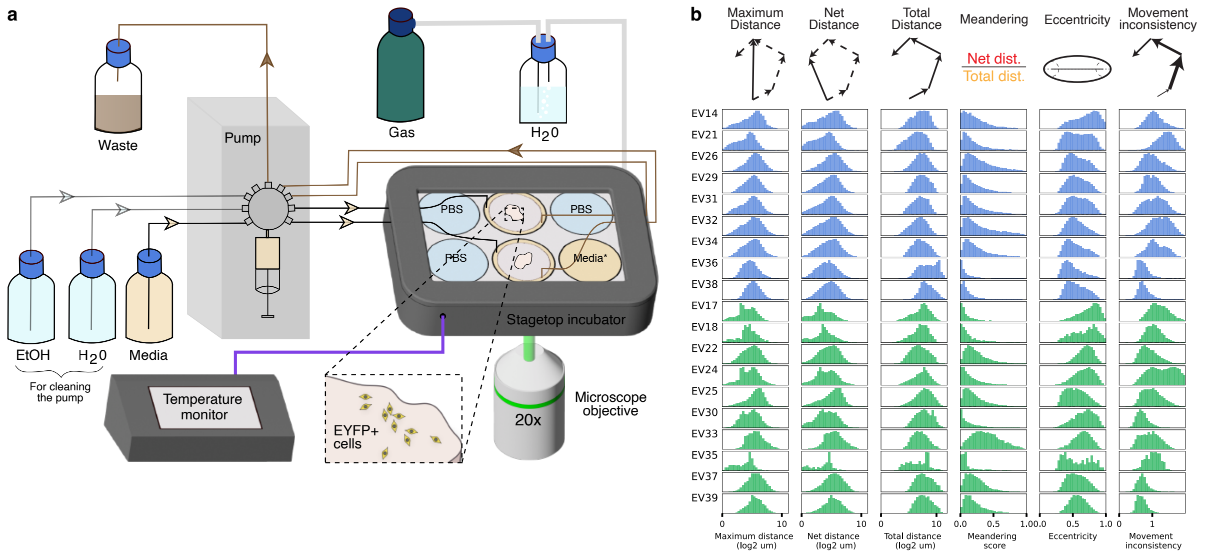


**Extended Data Fig. 3: Live imaging setup. a)** Schematic of the automated tissue perfusion set up. A programmable syringe pump with distributor head serves as a central controller for the partial replacement of culture media at an interval of 4 hours for up to a week at a time without any plate handling. **b)** Individual histograms for all live imaged samples displayed in Fig. 2d.

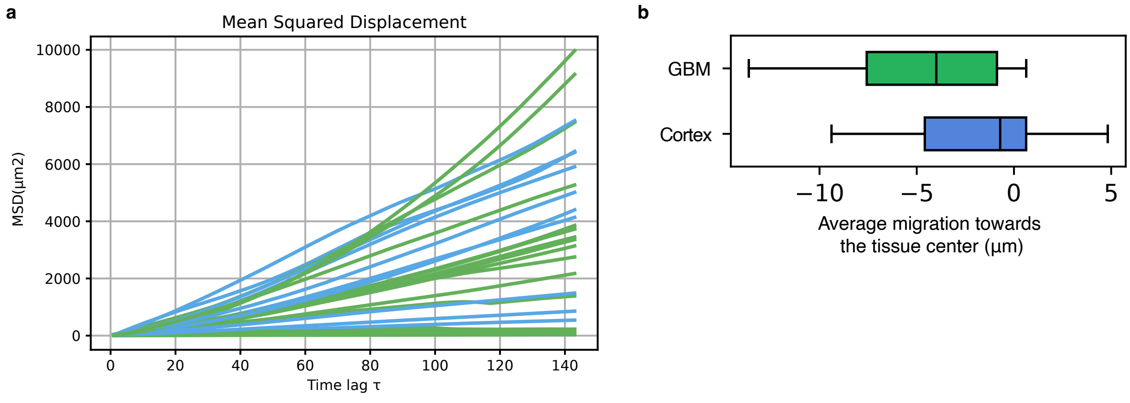


**Extended Data Fig. 4: Further descriptions of tissue growth dynamics. a)** Mean squared displacement of all cells across the first 3 days of imaging for each tissue sample. The colour delineates between glioblastoma and fetal cortex. **b)** Average migration of all cells in a tissue slice in relation to the tissue centre. Positive values represent movement toward the tissue centre while negative values represent movement away from the centre.

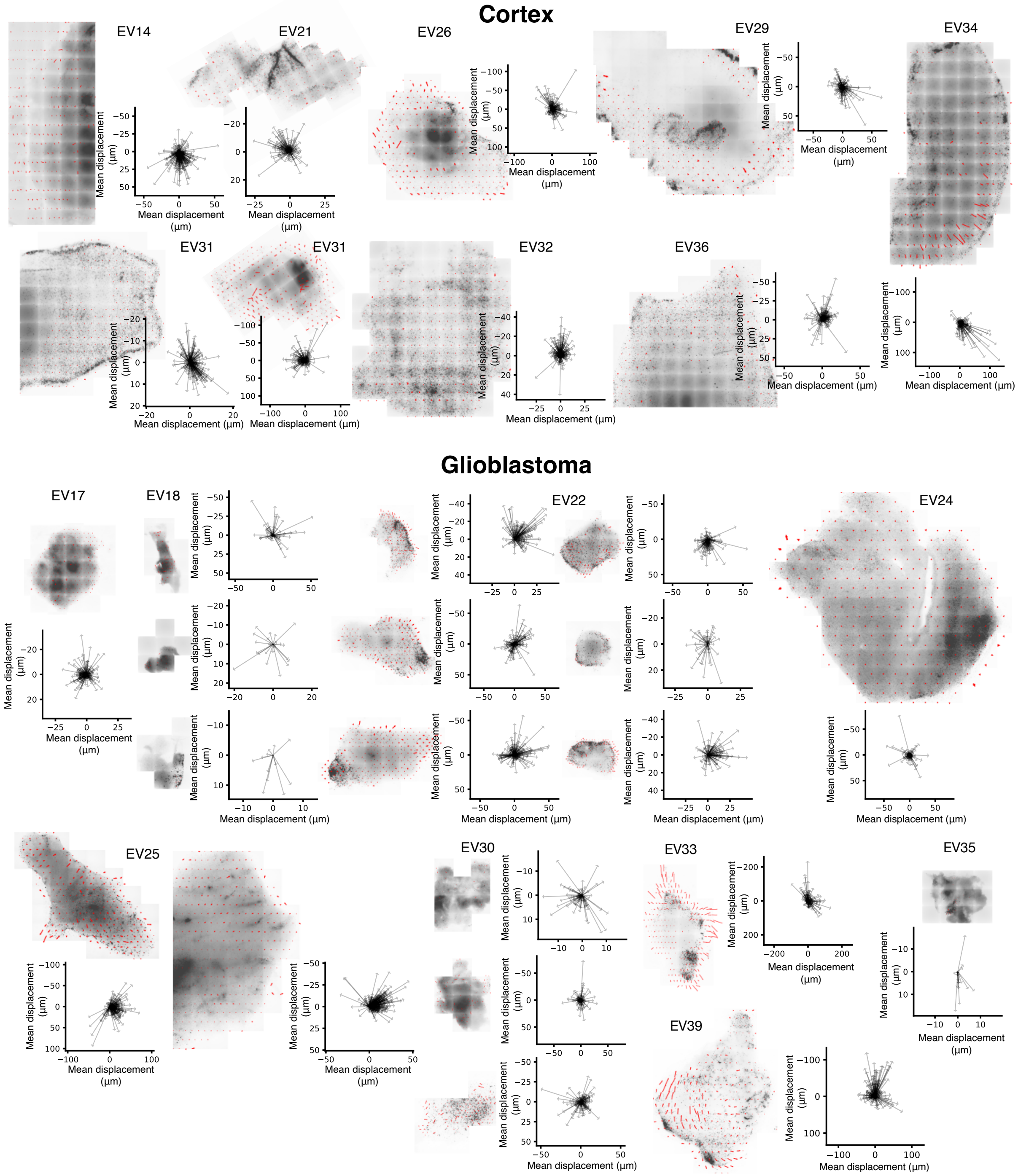


**Extended Data Fig. 5: Net movement of all fetal cortex and glioblastoma samples.** Wind plots correspond to adjacent fluorescence images (intensity inverted). In total 9 fetal cortex samples and 9 glioblastoma samples were included for analysis.

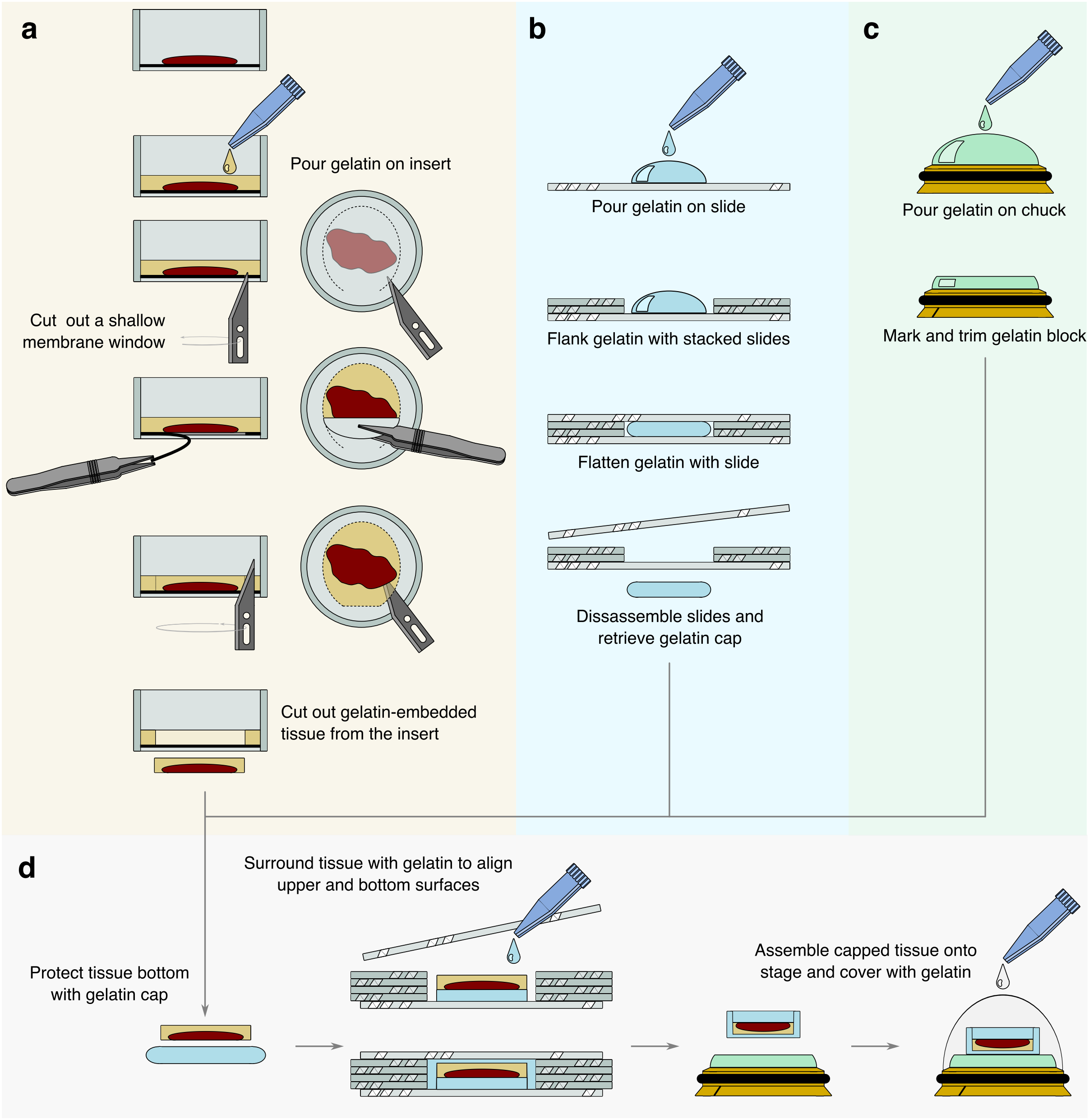


**Extended Data Fig. 6: Detailed protocol of resectioning.** Glioblastoma/fetal cortex sample-bearing inserts were retrieved post-live imaging and **a)** embedded in a gelatin (in 30% sucrose, 1X PBS) solution, after which the gelated block was exposed and cut out, then assembled together with a **b)** trimmed flat gelatin block and a **c)** cryosectioned gelatin stage atop a cryostat chuck. **d)** The final ensemble was covered in a protective layer of gelatin. Colours are illustrative.

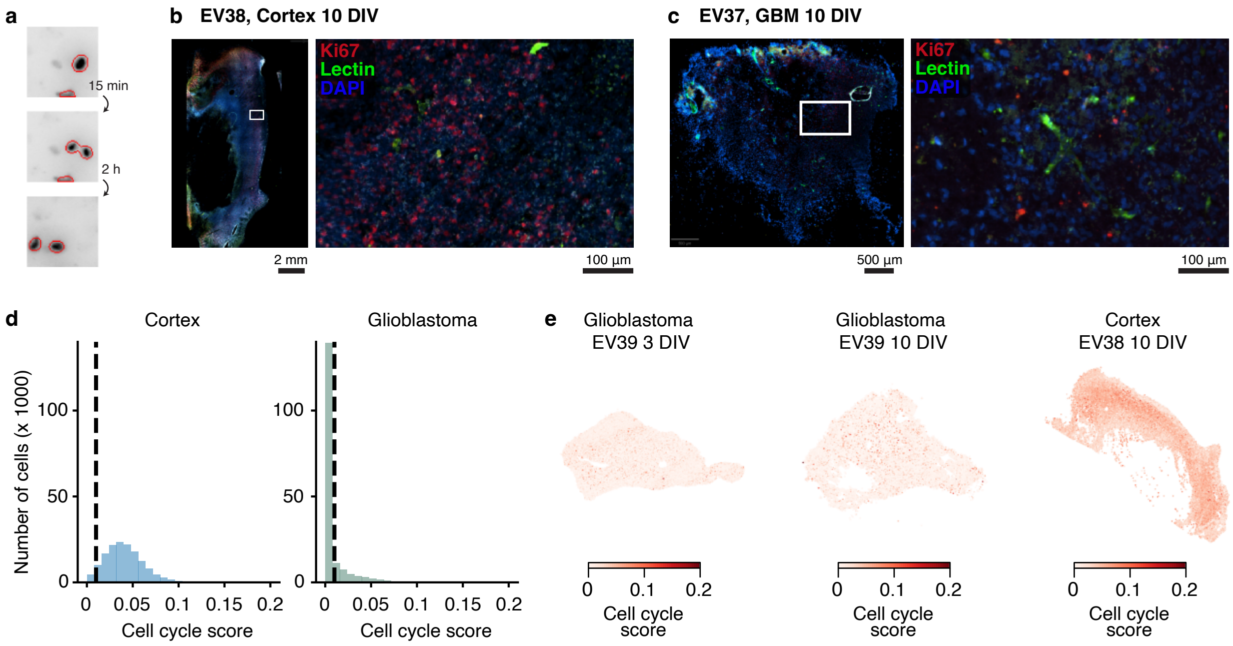


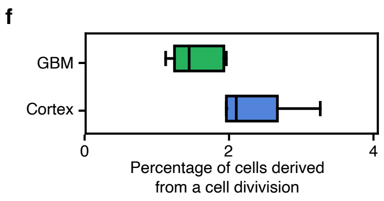


**Extended Data Fig. 7: Cell cycle and cell division. a)** Example of a cell division captured by timelapse imaging. **b)** Staining of cortex sample EV38 using anti-KI67, Lectin and DAPI after spatial transcriptomics. **c)** Identical staining for glioblastoma sample EV37 after spatial transcriptomics. **d)** Distribution of cell cycle scores of cortical and glioblastoma cells. Dashed line shows the 0.02 threshold used to identify cycling cells. **e)** Cell cycle scores plotted in a spatial context.

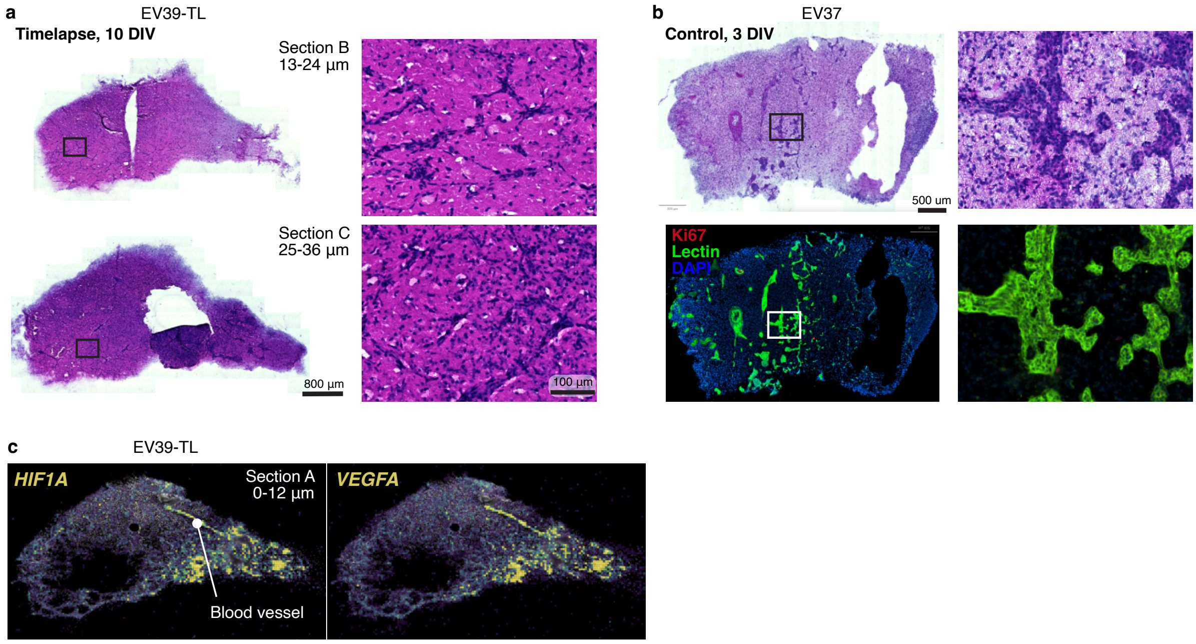


**Extended Data Fig. 8: Blood vessels. a)** Adjacent 10 days *in vitro* slice resectioned post-timelapse from 13 µm up to 36 µm from the PTFE membrane, stained with H&E. **b)** H&E and Ki67 and Lectin staining in EV37 glioblastoma sample post timelapse and spatial transcriptomics. **c)** Spatial transcriptomics of EV39 showing *HIF1A* and *VEGFA* expression.
